# Recurrent structural variation and recent turnover at the 17q21.31 locus in humans and great apes

**DOI:** 10.1101/2025.08.15.670618

**Authors:** Samvardhini Sridharan, Runyang Nicolas Lou, Scott Ferguson, Joana L Rocha, Rishi De-Kayne, Matthew W Mitchell, Alison N Killilia, Victor Borda, Santiago G. Medina-Munoz, Simon Gravel, CAAPA2 PopGen Working Group, Brenna Henn, Peter H Sudmant

## Abstract

The 17q21.31 locus in humans harbors several complex structural haplotypes including a ∼970kb inversion. Different inversion haplotypes have been associated with susceptibility to microdeletions causing Koolen-de Vries syndrome and variation in fecundity and recombination rates. Here, using 210 haplotype-resolved human genome assemblies and pangenome graph-based approaches we characterize 11 distinct structural haplotypes, several of which have not been previously described. Extending our analyses to a set of haplotype-resolved great-ape genomes, we characterize the structure of an independent inversion in chimpanzees which extends an additional 650kb, encompasses 5 additional genes, and is ∼2 million years younger than the human inversion. We further determine that gorillas exhibit an independent duplication of the *KANSL1* gene which may predispose them to Koolen-de Vries syndrome causing microdeletions. Using short read sequencing data we characterize 17q21.31 haplotype diversity worldwide in ∼5174 individuals from 107 populations finding increased frequencies of *KANSL1* duplication-containing haplotypes in both European and South Asian populations as well as 8 double recombination events between inverted and non-inverted haplotypes ranging in size from 20-180kb. Finally, using 626 ancient Eurasian human genomes we show the frequency of haplotypes containing *KANSL1* duplications has increased ∼6-fold over the past 12 thousand years in Europe. Together, our results highlight the dynamics, complexity, and recurrent, independent evolution of a medically relevant locus across humans and great apes.

## INTRODUCTION

Complex structural variants (SVs) play critical roles in human disease^1^, diversity^2,3^, and evolution^4,5^. Inversions are a particularly intriguing and understudied class of SV which have historically been challenging to assay as they are often flanked by large, complex, segmental duplications^1,6^. Nevertheless, across taxa inversions are powerful drivers of genetic and phenotypic diversity influencing traits such as behavior^7^, fitness^8,9^, and morphology^8,10^. Inversions can also act as barriers to gene flow between populations, promoting speciation and adaptation to different ecological niches and environmental pressures^11–13^.

One remarkable inversion in humans is found at the 17q21.31 locus spanning ∼970kb and impacting patterns of linkage disequilibrium over more than a megabase^14^. Haplotypes at this locus group into two major clades based on their inversion status: H1 haplotypes which are the most common in humans, and H2 haplotypes, which are inverted with respect to the H1 haplotype. Since its discovery, this locus has been associated with signatures of positive selection^14^, disease^15^, and recurrent structural rearrangement (“toggling”)^16^ throughout the great ape lineage, highlighting its complex evolutionary history.

The 17q21.31 locus is of particular medical importance because it harbors several genes associated with human disease. These include *MAPT*, which encodes the tau protein found in aggregates in Alzheimer’s dementias^17^, and *KANSL1*, disruption of which results in Koolen-de Vries Syndrome (KdVS)^15,18^. KdVS is characterized by developmental delay, intellectual disability, hypotonia, epilepsy, distinctive facial features, and congenital malformations across multiple organ systems^19^. While rare truncating mutations of *KANSL1* can cause KdVS^15^ it is more commonly caused by 17q21.31 microdeletions which occur at a frequency of ∼1/16,000 individuals. These microdeletions encompass 5 genes and are mediated by non-allelic homologous recombination (NAHR) between directly oriented segmental duplications which are present on the H2 (inverted) haplotype, but not the H1 haplotype. H2 carriers are thus at risk for KdVS. However, H2 haplotypes, which are at particularly high frequency in European populations, have also been associated with increased fecundity in some populations ^14,20^ and thus may be the target of positive selection^19^.

In addition to the non-inverted (H1) and inverted (H2) haplotypes, recent studies have identified additional structural complexity at the 17q21.31 locus^21,22^, including two independent partial duplications of the *KANSL1* gene on both the H1 and H2 haplotypes, as well as copy number variation of the *NSF* gene. Here, we set out to characterize the recent history and full extent of structural complexity at the 17q21.31 locus using both humans and non-human primate long-read haplotype-resolved genome assemblies, extensive human population diversity sequencing, and ancient genomes.

## RESULTS

### Resolving complex genome structures at the 17q21.31 locus with 210 haplotype-resolved genome assemblies

Recently generated long-read genomic resources provide an unprecedented opportunity to uncover structural diversity at complex loci such as the 17q21.31 locus. We collated 210 diverse haplotypes including the recently assembled T2T human genome to examine the structural complexity of this region. We first used PGRTK ^23^ (cite) to annotate the repeat architecture of these genomes (**Fig 1A, B, S1**). PGRTK identifies clusters of paralogous and homologous sequences within and between genomes enabling us to compare structures among haplotypes. Previous literature has established that alongside the major inversion haplotype groups H1 and H2, several additional regions of copy number variation exist on these haplotype backgrounds. These include distinct and independent partial duplications of the *KANSL1* gene on the H2 and H1 haplotypes, referred to as α and β respectively. Additionally partial duplications of the *NSF* gene, on either the H1 or H2 haplotype, are referred to as γ duplications.

**Figure 1.**
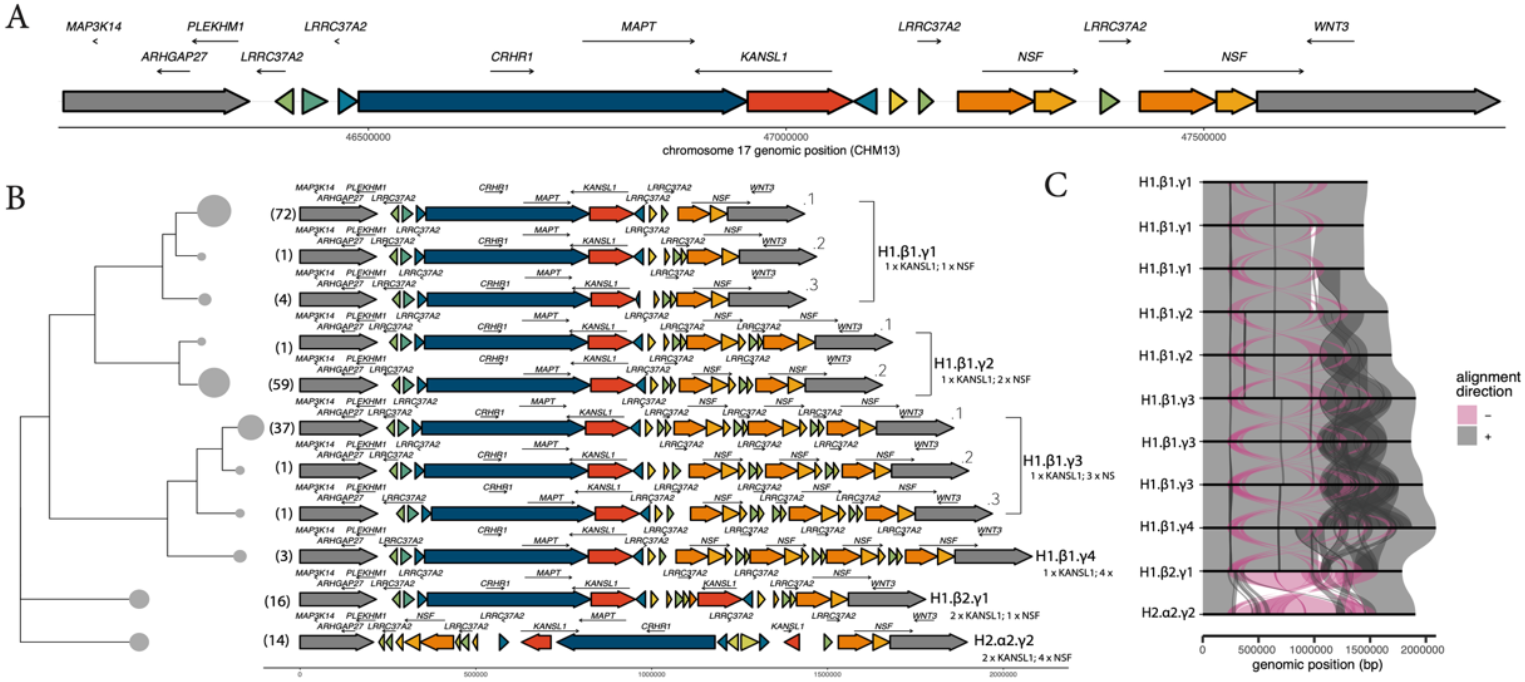
Distinct 17q21.31 structural haplotypes from human long-read genome assemblies: **a)** Schematic of haplotype structure at the 17q21.31 locus and flanking regions using PGRTK for the reference CHM13 genome. Large colored arrows represent principal bundles, which are individual units of repeats. Bundles of the same color represent homologous sequences. The directionality of the arrow represents the bundle orientation. Unique flanking regions are indicated in greyscale. Smaller labeled arrows denote genes and gene orientations. **b)** Schematic of 11 unique structural haplotypes found within 210 long-read haplotype-resolved assemblies. Each row represents a unique structural haplotype. Structural haplotypes are arranged using hierarchical clustering based on jaccard distance, with tip sizes scaled by the number of assemblies sharing each haplotype. **c)** Visualization of sequence alignment among 11 unique structural haplotypes. Each row represents a unique haplotype structure arranged in the same order as in **b**. Alignments are colored by orientation: grey denotes alignments between two structural haplotypes in the forward orientation and pink denotes alignment in the reverse direction.

Alongside our PGRTK-based classification of structures we used a pangenome-based approach (PGGB) ^24^ to cluster haplotype architectures (see methods) identifying a total of 24 unique structures (**Fig S1**). Many of these structures were highly similar differing only subtly in the composition of the segmentally duplicated *LRRC37A* gene. We thus further clustered structures based on their pairwise jaccard distance into 11 unique summary structures (**Methods**).

Structural differences between these haplotypes were confirmed by plotting a multiway structural alignment using SVbyEye (**Fig 1C**). These clusters exhibit finer resolution than previously developed nomenclature. For example, we identified three structures which can be classified as H1.β1.γ1, based on their single copies of the *KANSL1* and the *NSF* duplications, which we designate with additional numeric suffixes. We further identified two distinct H1.β1.γ2 haplotypes and three H1.β1.γ3 haplotypes. These distinct structures are mostly differentiated by copy number variation at the *LRRC37A2* gene, which flanks the inversion. Overall, the most commonly observed H1 structures were H1.β1.γ1.1 (37% of H1), H1.β1.γ2.2 (30% of H1), H1.β1.γ3.1 (19% of H1) and H1.β2.γ1 (8% of H1) - making up a total proportion of 94% of H1s (88% overall). Most of the variation in structural diversity we observed is located on H1.β1 haplotypes, which are the highest frequency. Only a single H1.β2 and H2.α2 structure were observed in our dataset and several previously described haplotype structures were not present in our long read assemblies (H1.β3.γ1, H2.α1.γ2, and H2.α2.γ1). However, these structures were observed at low frequency in our subsequent analyses of structural variation using short reads. Together, these data recapitulate previously described structures in higher detail and demonstrate additional complexity present in humans.

### Recurrent but distinct inversions in humans and chimpanzees

The recent completion of several haplotype-resolved T2T ape genomes provides the opportunity to contextualize the evolutionary history of this locus in great apes. To explore the population-level diversity in these non-human species we further sequenced and assembled two high-quality haplotype resolved gorilla genomes (four haplotypes) and assessed a set of recently assembled chimpanzee and bonobo genomes (Rocha et al. in prep). Previous work has demonstrated that the H2 orientation is the ancestral architecture of the locus. Intriguingly, fluorescent *in-situ* hybridization of a panel of several apes has suggested that the 17q21.31 inversion is segregating in orangutans, chimpanzees, and bonobos^16^ emerging independently in these species (i.e. inversion *toggling*).

We extracted the orthologous 17q21.31 sequence across five non-human great ape species and constructed structural alignments using SVbyEye (**Fig. 2A,B**). These alignments demonstrate the increasing complexity of segmental duplications in African great apes in comparison to orangutans, as has been previously described^16^. To understand how these structures relate to the human architecture we complemented these analyses by projecting human-identified PGRTK sequence blocks onto the great ape genomes (**Fig. 2C**). These projections lose resolution as a function of genetic distance particularly in rapidly evolving sequence such segmental duplications but still highlight major architectural features like the inversion and flanking regions.

**Figure 2.**
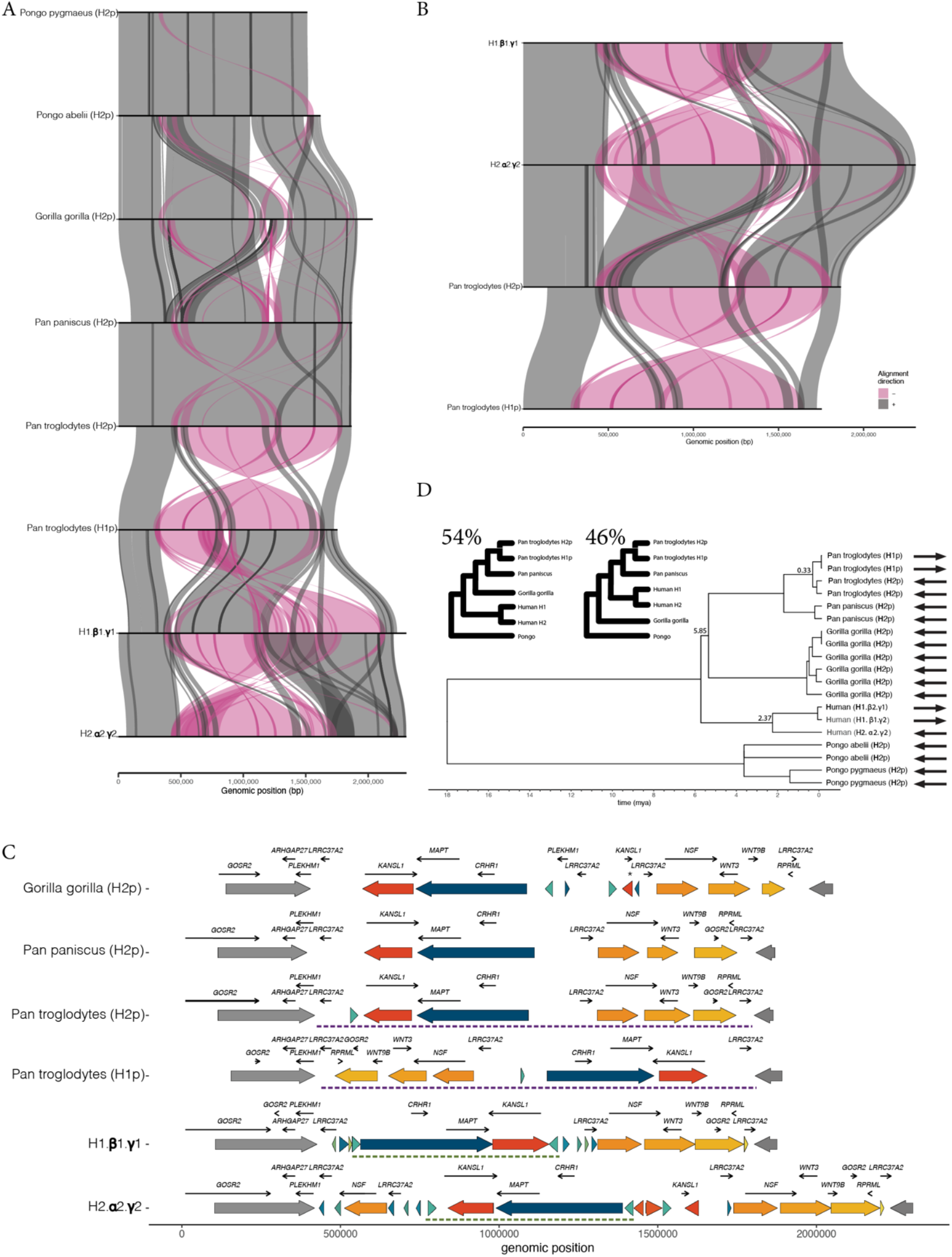
Characterizing the evolutionary context of the 17q21.31 inversion in the great ape lineage: **a)** Visualization of sequence alignments spanning the 17q21.31 locus and flanking regions in great apes. The sequences are extended from **Fig 1** by 2 Mb on both ends to characterize the shared homology of flanking regions across all species. Each row represents a representative haplotype from each of the great apes. Alignments are colored by orientation (gray = forward, pink = reverse). **b)** Alignment visualization comparing representative human haplotypes H1.β1.γ3 and H2.α2.γ2 and chimpanzee haplotypes H1p and H2p (as seen in **a**), reordered to highlight the distinct, and longer inversion found in chimpanzees. **c)** Unique structural haplotypes in great apes with two representative human structural haplotypes shown as reference. Human haplotypes are plotted along with gene labels and gene orientation. Sequences are visualized with PGRTK trained on human structures with extended sequence as seen in **a**. Unique flanking regions are shown in dark grey and lighter grey. Colors are broadly consistent with **Fig 1b** although the longer genetic interval used here results in slight differences. **d)** Dating the divergence of great ape and human 17q21.31 haplotypes. Trees were inferred using a large non-duplicated sequence located within the human inversion (i.e. the blue bundle in **c, see methods)**. Several important split times are annotated in the tree. The inset shows two main topologies that are observed via bootstrapping and their frequencies.

Our analyses revealed several striking, lineage specific structural features at this locus. In gorillas we identified a 500kb duplication overlapping the *KANSL1* gene. This is an independent duplication of the gene distinct from that observed in human haplotypes. This duplication corresponds to an annotated gene sequence in the recent T2T gorilla genome assembly, though it is unclear if it is a functional duplication. Furthermore, these *KANSL1* duplications are in direct orientation predisposing these haplotypes to NAHR-driven microdeletions/duplications. Thus, gorillas may be at risk for Koolen-de-vries syndrome. We also identified two inverted chimpanzee haplotypes. These inversions are in the same orientation as the human H1 haplotype. We refer to this haplotype as the H1p haplotype, and the ancestral primate haplotype as H2p. The chimpanzee H1p/H2p inversion spans ∼1.4Mbp (**Fig 2B, S2**) encompassing the entirety of the human inversion and further extending an additional ∼650kb overlapping several genes including *WNT3, WNT9B, GOSR2*, and *RPML* (**Fig 2B**). The human and chimpanzee inversions share one approximate breakpoint though the other is distinct (**Fig S2**). Thus, not only is the chimpanzee inversion the result of an independent event, but it is a completely distinct structural variant. This provides further evidence that the H2 orientation is the ancestral structure in humans and chimpanzees, as while the H1 and H1p haplotypes differ structurally by hundreds of kb the H2 and H2p haplotypes exhibit highly similar structures.

We next sought to compare the timing of the different inversion events by aligning ∼400kbp of unique (non-duplicated) sequence within the shared inversion block across humans and great apes (**Fig. 2D, methods**). The human H1/H2 inversion exhibits a coalescence time of approximately to 2.38 million years ago, confirming a relatively deep coalescence of the two haplotypes^21^. The chimpanzee H1p/H2p inversion however emerged much more recently ∼330,000 years ago. These analyses also revealed extensive incomplete lineage sorting across humans, gorillas, chimpanzees, and bonobos. 54% of trees exhibited a topology with humans as an outgroup to the *pan* and gorilla lineages with the canonical topology represented 46% of the time. Together, these results demonstrate that the 17q21.31 locus is highly dynamic across great apes with distinct but overlapping structural variants occurring independently in several taxa over vastly different timescales.

### Timing and recurrence of human 17q21.31 structural haplotypes

Long-read haplotype-resolved assemblies phase genome structures with linked SNPs which can be used to trace the evolutionary history of complex genome architectures. Extended suppression of recombination between the H1 and H2 haplotypes enables dating of the coalescent time of the inversion to ∼2.38 million years. However, recombination between H1 haplotypes will disrupt patterns of linkage disequilibrium (LD) making it challenging to assess the emergence of specific H1 substructures. We sought to identify high LD blocks adjacent to the distal region of H1 structural variation (containing the *KANSL1* and *NSF* duplications). Such regions can be used to trace the history of these structures. We identified 100kb of sequence adjacent to the structurally variable portion of H1 with a consistent tree topology indicating high LD and reduced recombination (**Fig 3A, B**). Using this sequence we constructed a coalescent tree across 210 human haplotypes with the T2T chimpanzee genome as the outgroup. This tree reveals that H1 *KANSL1* duplications (i.e. H1.β2) are largely found in one cluster with a handful of samples falling in an independent region of the tree, potentially resulting from an independent origin, or a recombination event.

**Figure 3.**
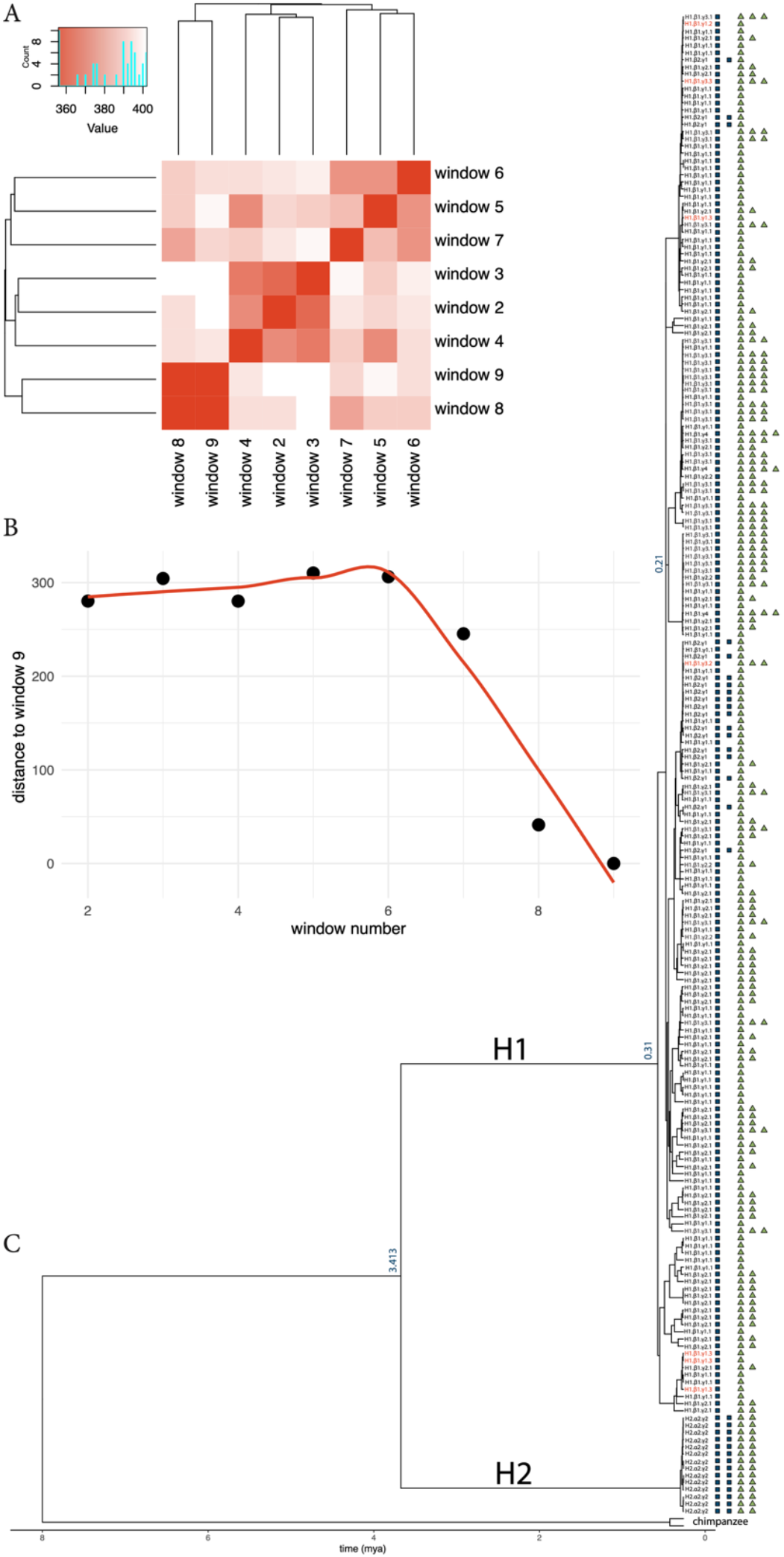
Timing and recurrence of human 17q21.31 structural haplotypes: **a)** Robinson-Foulds (RF) distance clustering heatmap across phylogenetic trees constructed with non-overapping 50 kb windows across all human long-read assemblies. Windows are extracted from a large non-duplicated sequence located within the human inversion (i.e. the blue bundle in **Fig 1)**. Darker red indicates greater similarity (lower RF distance) between two trees. Dendogram summarizes the topological similarity across all windows. Inset histogram displays the distribution of RF distance values across all comparisons. **b)** Euclidean distance along PC1 and PC2 axes between the phylogenetic tree constructed at window 9, the closest unique sequence to *KANSL1*, and trees at all other windows. A loess-smoothed curve is shown in red. **c)** Dated maximum-likelihood tree constructed using sequences at windows 8 and 9 and rooted with T2T chimpanzee haplotypes as the outgroup. Human-chimpanzee split-time was used to date the tree. Select internal nodes are labeled with divergence time. Tip symbols indicate copy-number variation of *KANSL1* (blue squares) and *NSF* (green triangles).

In contrast, *NSF* duplications (γ2, γ3) likely exhibit several independent origins throughout the tree. One particularly large cluster of γ3 duplications was identified, within which all observed γ4 duplications were also found. We also find independent origins of the sub-haplotypes cataloged in **Fig 1A** including multiple origins of the H1.β1.γ1.3 structure. Together, these results highlight that the structure of H1 haplotypes at the 17q21.31 locus is highly mutable and predisposed to recurrent structural variation. This includes large-scale gene duplications such as *KANSL1* and *NSF* as well as smaller SVs including copy number changes of *LRRC37A*.

### Population diversity of the 17q21.31 locus worldwide using short read data

Despite their immense power, existing long-read datasets remain limited in sample size and population diversity. Nevertheless, large-scale short-read sequencing datasets from diverse human populations have recently been generated. To characterize patterns of 17q21.31 variation across the world we analyzed short-read sequencing data from 5174 samples encompassing 107 populations representative of human diversity worldwide (see **Methods, Figs S3-5**). Using SNPs tagging the inversion as well as read-depth based copy number quantification we estimated frequencies of four simplified haplotype architectures which could be accurately determined from the short read data: H1.β1, representing H1 haplotypes without *KANSL1* duplications; H1.β2+, representing H1 haplotypes with *KANSL1* duplications; H2.α1 representing H2 haplotypes without *KANSL1* duplications; and H2.α2+ representing H2 haplotypes with *KANSL1* duplications (**Fig 4A-D**).

**Figure 4.**
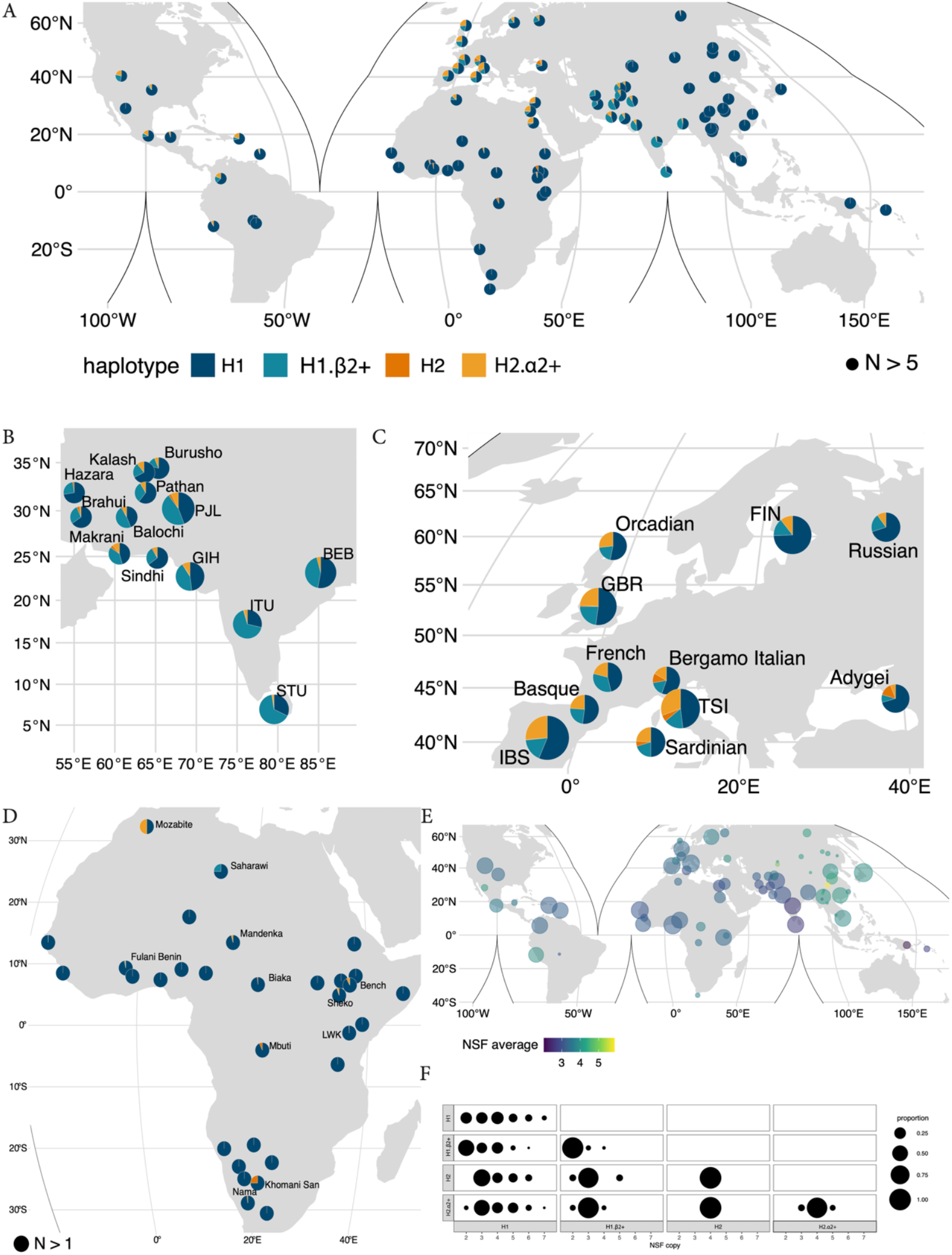
Population diversity of the 17q21.31 locus worldwide using short read data: **a)** World map indicating the frequency of direct H1.β1, H1.β2+ and inverted H2.α1 and H2.α2+ 17q21.31 haplotypes in 4585 individuals from 193 populations. Insets show the frequency distribution in South Asia (**b)**, Europe (**c**) and Africa (**d**). Pies are scaled by sample size in **b** and **c**. Only populations with five or more samples are plotted in **a-c**, whereas **d** shows populations with as few as a single sample. In **d**, only populations with H1.β1 frequencies less than 1 are labeled. See **Supplementary Table 2** for exact sample sizes and haplotype frequencies of all populations. **d)** World map indicating the average diploid *NSF* copy number in 3352 individuals from 147 populations. Points are scaled based on sample size, with only populations with five or more samples plotted. See **Supplementary Table 3** for the exact sample sizes and average diploid copy numbers of all populations. **e)** The distribution of *NSF* copy numbers in individuals with different inversion and *KANSL1* duplication genotypes.

Worldwide, the most common haplotype is H1.β1,confirming previous work using more limited genome sequencing datasets^21,22^. However, we observe extreme population stratification of complex haplotypes at this locus. European populations exhibit the highest worldwide frequency of H2.α2+ haplotypes (10%-30.58%) and intermediate haplotypes frequencies (10.9%-31.5%) of H1.β2+. In contrast, South Asian populations, which have been historically underrepresented in sequencing datasets, exhibit the highest worldwide frequency of H1.β2+ haplotypes (14.6%-66.8%) and lower levels of H2.α2+ (1.4%-10.9%). East Asian populations are essentially fixed for H1.β1 haplotypes. We also assessed whole genome sequencing data from 368 African genomes from the CAAPA2 consortium demonstrating limited haplotype diversity in Africa, with most populations exhibiting H1.β1. However, we observed H2.α1 haplotypes at low frequency in Khomani San, Nama, Mbuti, Biaka, Bench, and Sheko populations. H2.α2+ haplotypes were also present at low frequencies in some African populations. These results confirm previous analyses highlighting the presence of rare H2.α1 haplotypes almost exclusively in African populations^21^ however also highlight that *KANSL1* duplication-containing haplotypes (including the inverted H2.α2+ haplotypes) have increased dramatically in both Europe and South Asia.

While short-read sequencing data do not enable characterization of phased *NSF* duplications (i.e. γ2+) we calculated diploid *NSF* copy number genotype frequencies worldwide confirming the highest *NSF* copy numbers in East Asia (**Fig 4E**). We find that the overwhelming majority of *NSF* duplications occur on the H1.β1 background. However, we do observe evidence of both *NSF* duplications and deletions happening at low frequencies on H2.α2 background (**Fig 4F**).

Together, these results provide the most comprehensive catalog of 17q21.31 diversity in humans to date and highlight the extreme stratification of haplotypes between continental populations.

### Double recombinants and complex gene conversion events between structural haplotypes

Inversions can strongly suppress recombination impacting patterns of linkage disequilibrium (LD), however genetic exchange between inversion alleles can occasionally occur through double recombination or gene conversion events. To understand how the 17q21.31 inversion has impacted patterns of recombination we used SNPRelate^25^, a windowed PCA-based analysis, to cluster 3135 short-read samples in 10kb sliding windows across the locus (**Fig 5A, B**). Most individuals consistently clustered into one of three groups corresponding to H1/H1, H1/H2, H2/H2 genotypes throughout the non-duplicated part of the inversion and even ∼80kb beyond the proximal inversion breakpoint, highlighting the far-reaching effect of inversions in suppressing recombination. Despite the persistence of this long 900kb LD block, we identified at least three instances of recombination at the proximal end of the extended LD block (**Fig 5C**) occurring primarily in African individuals (32/56, 57%). We also noted several individuals exhibiting 20-180 kb stretches showing switching between genotype groups within the inversion region, indicating double crossover recombination events, or possibly, long stretches of interlocus gene conversion (**Fig 5A-C**). Of the 8 identified switch events, two exceeded 140 kb, occurring in a single European individual and a single South Asian individual respectively, and represent cases of double crossover recombination. The remaining events range from 20-70 kb and were found almost exclusively in African populations (51/54, 94%). We also noted that this windowed PCA approach identified more complex clustering patterns at the proximal and distal ends of the inversion locus that contain duplications (**Fig 5B**). We found that these patterns corresponded strongly with the complex structural haplotypes, thus suggesting that complex structural variation can be detected from short read-based SNPsin some cases. Together these results demonstrate that despite extensive inversion-driven recombination suppression, substantial exchange has happened between vastly divergent haplotypes.

**Figure 5.**
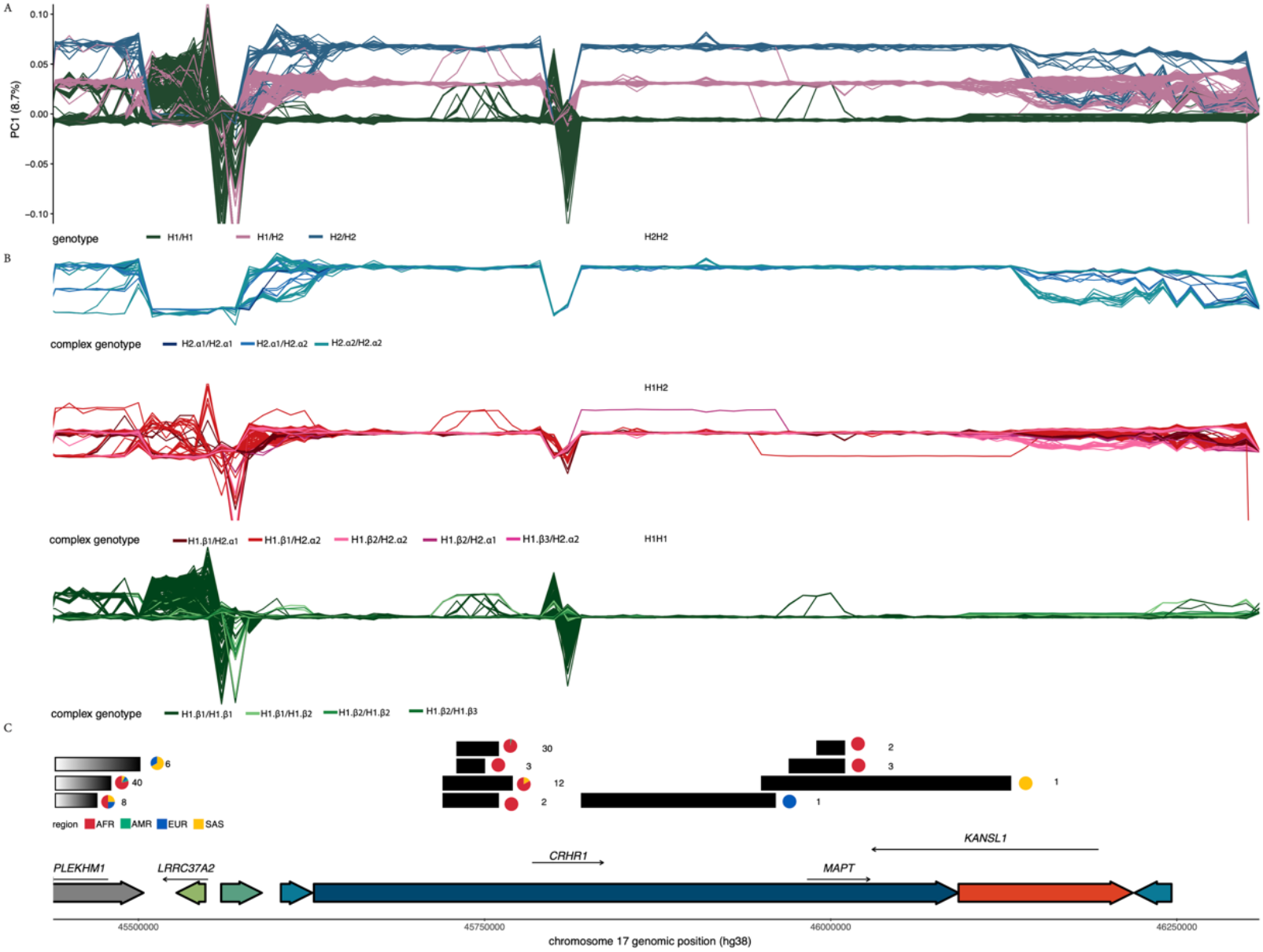
Population specific instances of recombination events at the 17q21.31 locus: The x-axis represents genomic position across a subset of the 17q21.31 locus including the inversion region and surrounding flanking sequences. A PGRTK-based haplotype structure of the reference genome (hg38) is shown below the figure. **a)** 10kb sliding window PCA. Each colored line represents the variation in the position of an individual on the first principal component axis, and is colored by inversion genotype. **b)** 10kb sliding window PCA faceted by inversion status and colored by complex genotype. **c)** All recombination events identified in the region of interest. Gradient blocks represent single recombination events wherein the right edge of the block represents the recombination breakpoint. Black boxes represent double recombination events. Numbers next to each block indicate the number of individuals with that recombination event, and pie charts indicate the population of these individuals.

### Ancient genomes track the rise in frequency of *KANSL1* duplications in Europe

To determine how 17q21.31 structural haplotype frequencies have changed in Europeans over recent history we analyzed 626 ancient genomes spanning ∼10 thousand years of human history^26^ **(Fig 6A)**. European Hunter Gatherer populations, dating from ∼5000-10,000 years before present (BP) exhibited extremely low haplotype diversity with H1.β1 making up >95% of haplotypes **(Fig 6B)**. Higher frequencies of H2.α2+ haplotypes appear ∼7 to ∼5 kya BP in Early and Neolithic Farmers (25%-31%). Steppe Pastoralists also exhibit H2.α2+ haplotypes (10%) as well as high frequencies of the H1.β2+ haplotype (30%). To understand the temporal dynamics of haplotype frequency changes over the last 10 kya we fit a multinomial logistic regression model to these ancient genomic derived data **(Fig 6C)**. This model demonstrates that while H1.β1 was the dominant haplotype ∼11 thousand years ago with a frequency of >90%, it has subsequently declined ∼1.6-fold. Concomitantly, the H1.β2+ and H2.α2+ haplotypes have increased from ∼1% and ∼7% to 29% and 21% respectively. Together, these data demonstrate that 17q21.31 haplotype diversity in Europe today is the result of a massive increase in frequencies of the *KANSL1* duplication in both H1 and H2 haplotypes over recent history.

**Figure 6.**
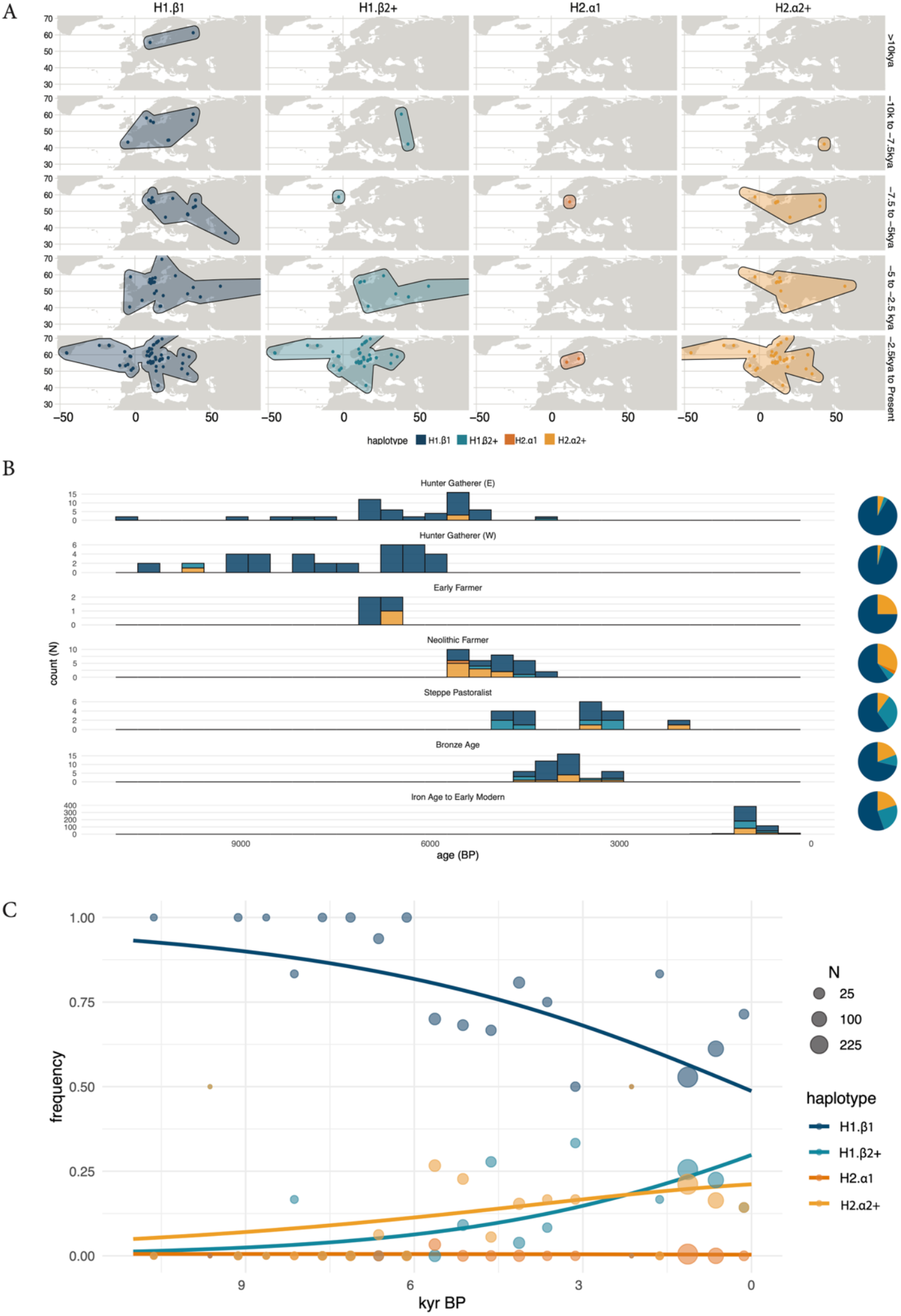
Ancient genomes track the rise in frequency of *KANSL1* duplication in Europe: **a)** Geographic distribution of historical samples across Western Eurasia shown in 2.5 kya time slices from 10 kya to the present. Samples are split based on their haplotype structure (H1.β1, H1.β2+, H2.α1, and H2.α2+) and age in each subplot, with dots marking individual sampling locations. **b)** Temporal distribution of 17q21.31 haplotypes across historical West Eurasian populations. Stacked bar plots show the number of haplotypes present in each population over a time bin. Time is shown on the x-axis in years before present (BP). Pie charts on the right summarize the overall haplotype frequency for each population, with the. **c)** Modeled temporal trajectories of 17q21.31 haplotype frequencies over the past 10 ky using a multinomial logistic regression model. Each point represents the observed haplotype frequency in a time bin (all the populations are pooled), with point size scaled by sample size.

## DISCUSSION

Here we leverage long-read haplotype-resolved human genomes, newly assembled T2T and population-scale long-read sequenced great ape genomes, diverse human short-read data, and ancient DNA to assess the diversity and recent evolution of the 17q21.31 locus. Together, our results provide the most comprehensive assessment to date of this remarkable region of the genome providing new insights and clarifying previous works. For instance, while it has been established that the H2 haplotype is the ancestral orientation of the 17q21.31 locus, the presence of the inversions in chimpanzees, bonobos, and orangutans has been interpreted as recurrent “toggling” of the inversion across great apes. Here, by using population-scale haplotype resolved assemblies from chimpanzees we resolve the structure and sequence of the chimpanzee 17q21.31 inversion. We find that the chimpanzee inversion is substantially larger than the human inversion. While it overlaps the human inversion across ∼750kb of sequence it extends an additional ∼650kb. Furthermore, we find that this inversion occurred far more recently than the human inversion, ∼330,000 years ago, in contrast to previous estimates^16^. Thus, “highly mutable and recurrent” might be a more apt description of the inversion locus than “toggling” as independent inversion events with distinct breakpoints have occurred on different lineages. However, additional population-scale long-read sequencing of primate genomes will be required to determine the sequence and structures of the bonobo and orangutan inversions and discern their evolutionary history.

The H2 haplotype has been associated with increased fecundity in Icelandic^14^ and Polish^20^ populations and has thus been suggested to be the target of selection in Europeans. Indeed, H2 haplotype frequencies are highest in European populations. We used ancient genomes to reconstruct allele frequency trajectories of 17q21.31 haplotypes finding that while H1.β1 haplotypes have declined in frequency markedly over the past 10,000 years, H1.β2+ and H2.α2+ haplotypes have increased in frequency substantially: ∼29-fold for H1.β2+ and 3-fold for H2.α2+. Thus, the group of haplotypes containing duplications of the *KANSL1* gene has increased in frequency ∼6-fold. Prima facie such an increase in frequency would correspond to a selection coefficient of approximately 0.015, similar to the average selection coefficients for regions such as lactase persistence. Critically, however, this estimate is an extremely crude oversimplification which neglects the small sample sizes in the ancient data and the importance of modelling the complex demographic history of Europe. Nevertheless, these results provide an intriguing alternate hypothesis: that duplications of the *KANSL1* gene may be the targets of selection as opposed to the inversion itself.

The availability of increasingly diverse population-scale short-read sequencing of humans enabled us to characterize 17q21.31 haplotype diversity worldwide. Our results confirm previous work highlighting the prevalence of the H1.β1 haplotype in Africa with the ancestral H2.α1 haplotypes present at low frequencies in some populations such as the Khomani San. However, our analyses also provide some of the first observations of 17q21.31 diversity in South Asian populations, which have historically been undersampled. We find surprisingly high frequencies of the H1.β2+ and H2.α2+ haplotypes across South Asia. These results mirror those of Europe with the two *KANSL1* duplication-containing haplotypes showing increased frequencies compared to African and East Asian populations, albeit with differing relative frequencies of the H1.β2+ and H2.α2+. These results potentially suggest that a recent increase in duplicated haplotypes may have also occurred in South Asia mirroring the increase in Europe, though additional analyses are needed to confirm that hypothesis.

Finally, we discovered several double recombination events inside the 17q21.31 inversion locus. Intriguingly, these events are overwhelmingly found in African populations, many of which have very low H2 haplotype frequencies. Therefore, these double recombination events are likely quite old, having occurred when H2 haplotypes were at higher frequencies in Africa. We also identified two other much longer double recombination events present in two non-African individuals. In addition, we identified a region intersecting the *CRHR1* gene where H1 and H2 haplotypes are completely homogenized and individuals cannot be assigned to one haplotype or the other (**Fig 5A-C**). This finding has previously been reported and explained as the result of an ancient double recombination event^21^. Crucially, unlike the other double recombinants that we identified, this event has likely been fixed in H2 haplotype, and the original sequence associated with the H2 haplotype has been lost in all human populations. Except for these rare double recombination events, however, we found that recombination is completely suppressed across the inversion. An additional consequence of segregating inversions is that by suppressing recombination they can promote the accumulation of deleterious variation^9^. It remains an unanswered question how the increased frequency of 17q21.31 inversion in specific human populations may affect mutational load.

The importance of the 17q21.31 locus in human disease, human diversity, and evolution has been the focus of many studies since its discovery. Nevertheless, long read sequencing, ancient genomes, and additional human population datasets are continuing to increase our understanding of this region. Our insights synthesize previous work with these new advances in genetics and genomics highlighting both new insights and posing new avenues for future research.

## METHODS

### Datasets

Short-read sequencing data were compiled from high-coverage resequencing of 5174 individuals from the 1,000 Genomes Project (1KG), the Human Genome Diversity Panel (HGDP) and the Simons Genome Diversity Panel (SGDP). We also obtained unpublished short-read sequencing data of 318 individuals from the CAAPA2 Data consortium. In total we were able to infer the complex genotypes of 4585 individuals with high confidence from these contemporary samples. Among these 697 are trios from 1KG and the remainder 3888 are unrelated individual samples compiled from 1KG, HGDP, SGDP, and CAAPA2. The trios are only used for method validation purposes. Genotypes from 626 historical samples from between -12kya and 1000 ya were obtained from ^27^. Historical samples were assigned to populations based on their original publications ^5,26^. 216 long-read haplotype assemblies were compiled from the HPRC and HGSVC ^3^. In addition we included the GRCh38 genome and CHM13 reference genomes to our dataset^28^. Haplotype assemblies were only included if they spanned the 17q21.31 locus. This excluded three assemblies `HG0015#hap1, HG00320#hap2, and NA21093#hap1`from our dataset, although they were used in characterization of complex structures in the locus. These three assemblies that were excluded appear to have been misassembled across the locus as they were either discontiguous, or had a partial *KANSL1* duplication that could not be cross-validated with short-read sequencing data. We obtained great ape genome assemblies for orangutans, gorillas, bonobos, and chimpanzees from three sources: GenomeArk (eight haplotypes, two per species), and 57 newly generated and unpublished gorilla, chimpanzee, and bonobo genomes. For the purposes of this study, we include the unpublished assemblies for the 17q21.31 region, with full genome releases planned alongside separate publications currently in preparation.

### Characterization of complex structures at the 17q21.31 locus in humans

We extracted the 17q21.31 region including flanking sequences from the GRCh38 reference genome (chr17: 45275488-46921902). We then used minimap2^29^ (-x asm5) to identify the homologous sequences in all the samples in the HGSVC and HPRC genomes. We performed a principal bundle decomposition using PanGenomic Research Tool Kit (PGR-TK) for 216 of the HGSVC and HPRC genome assemblies with the parameters of -w 80 -r 12 --min-span 32--bundle-length-cutoff 10000 --min-branch-size 16 --min-cov 0^23^. We examined the results, and excluded five assemblies that were potentially misassembled. We further excluded bundles that were unique to just one haplotype, or those whose length were in the lowest quantile across all the remaining bundles (i.e. < 23kb). We grouped the haplotypes into representative structures based on the order and orientation of the principal bundles. The position of genes along haplotypes was determined by mapping coding sequences to haplotypes using minimap2^29^ `(-c -x asm5)`. We chose one haplotype within each unique structure and used PGGB to build a pangenome graph^24^. We then used odgi similarity to generate a matrix of jaccard distance between each unique haplotype structure and made a neighbor joining tree to cluster these haplotype structures. We ran minimap2^29^ `(-x asm5 -c -- eqx -D -P --dual=no)` on pairs of assemblies to create sequence alignments, which were then inputted to SVByEye^30^for visualization.

### Characterization of complex structures at the 17q21.31 locus in great apes

We used minimap2^29^ (-x asm20) to identify the orthologous sequences in the great ape assemblies to the human 17q21.31 locus in GRCh38 reference genome (chr17:45275488-46921902). Because we observed an inversion in chimpanzees that spans a longer region than the inversion in humans, we then manually extended these sequences by 2Mb on either flank in both great apes and all human haplotypes, so that these sequences span the entirety of the chimpanzee inversion. We then reran PGRTK on these extended human sequences with the same settings and projected the principal bundles to the great ape sequences. Within each species, we categorized these haplotypes by their inversion, *KANSL1* and *NSF* duplication status. We then selected the one representative haplotype for each unique structure in each species for visualization. We ran minimap2^29^ and SVByEye^30^ on pairs of great ape and human haplotypes with the same settings except -x asm20 was used as a flag in minimap2^29^ instead of -asm5. To estimate the divergence time of structural haplotypes across great apes, we first identified the largest region of shared homology within the inverted segment (corresponding to the blue bundle in Fig. 1A). We performed pairwise alignments using minimap2^29^, with the CHM13 human reference sequence as the query, against 17 great ape assemblies: all great ape haplotypes from GenomeArk (including 4 orangutan, 2 gorilla, 2 bonobo, and 2 chimpanzee haplotypes), 4 additional gorilla haplotypes, the inverted chimpanzee haplotype, and representative human haplotypes H1.β1.γ1, H1.β2.γ1, and H2.α2.γ2. Multiple sequence alignment was performed using kalign^31^. Phylogenetic trees were inferred using IQ-TREE with the Jukes-Cantor substitution model and 1000 bootstrap replicates Divergence times were estimated using the orangutan–human/chimpanzee/gorilla split 18.13 mya as a calibration point^32^. Trees were visualized using the ggtree R package^33^.

### Evolutionary history of structural variation at the 17q21.31 locus in humans

To date the emergence of different H1 haplotype structures we first needed to find a unique sequence at the 17q21.31 locus that is physically linked to the *KANSL1* and *NSF* duplications and not broken down by recombination. The largest region of shared homology within the inverted segment (corresponding to the blue bundle in Fig. 1A) in CHM13 is also the closest in proximity to the *KANSL1* duplications. Therefore, we used similarities in phylogenetic tree topology as a proxy for linkage disequilibrium across this region.This region was divided into nine non-overlapping 50 kb windows using seqkit (sliding -s 50000 -W 50000)^34^. All 210 human haplotypes that were included in previous analyses were included to construct the trees, as well as 2 GenomeArk chimpanzee haplotypes. For each of the nine windows, sequences were aligned using kalign^31^, and maximum likelihood trees were inferred with IQ-TREE^35^ using the Jukes–Cantor model and 1000 ultrafast bootstrap replicates `(-m JC -bb 1000 -nt AUTO)`. Trees were rooted using the two chimpanzee haplotypes as an outgroup. To quantify topological similarity between trees, we computed pairwise Robinson–Foulds (RF) distances. An RF distance matrix was then constructed for all pairwise tree comparisons, and the windows with the lowest distance to the right-most window (i.e. the window that is closest to the *KANSL1* duplication) were selected. We then merged the selected windows, and reran IQ-TREE using the chimpanzee-human split time 7.74 mya as a calibration point for the emergence of haplotype structures^35^.

### Using contemporary short read genomes to understand 17q21.31 population diversity in modern human populations

#### Tag SNP based inversion genotyping

In order to determine the genotype status of individuals in our dataset, we used a total of 1271 tag SNPs to genotype the 17q21.31 locus based on the method used in^27^ For each individual, we constructed a genotype vector by coding observed genotypes at these SNPs as 0 (homozygous reference), 1 (heterozygous), or 2 (homozygous alternate). Then, Euclidean distance was calculated between the observed genotype vector and expected genotype vectors for H1/H1, H1/H2, and H2/H2. Each individual was assigned the inversion genotype with the minimum Euclidean distance to their observed genotype.

#### Read-depth based copy number genotyping at NSF and KANSL1

Copy number genotypes for the 1000 Genomes, HGDP, and SGDP, and CAAPA2 individuals were estimated using read depth as described in Bolognini et al^5^. In brief, read depths were quantified from CRAM (CAAPA2) or BAM (1000G, HGDP, SGDP) files. CRAM files from CAAPA2 were based on a different version of the GRCh38 reference genome, and to ensure consistency across datasets, we remapped this data to the same version of the reference genome as the other three datasets (GRCh38_full_analysis_set_plus_decoy_hla). We generated d4 files from these BAM files. From the d4 files we calculated the average read depths in 1,000 bp sliding windows in 100 bp steps across the 17q21.31 locus. These depths were normalized to the average read depth within a control region chr17:42800000-46000000 within the 17q21.31 locus for each individual, in which no copy number variation was observed in more than 3,000 individuals (i.e. individuals from the 1KGP and HGDP samples). To determine the copy number of the α and β segments of the *KANSL1* duplication, we averaged the copy number of windows that overlapped with the unique region of each duplication (α: chr17:46143000-46238000 and β: chr17:46095000-46123000). In the case of *NSF* copy number, since the GRCh38 reference genome has two copies of *NSF*, we summed the normalized read depth across two target regions corresponding to the two *NSF* copies (chr17:46,336,376–46,489,410 and chr17:46,564,311–46,707,123). CAAPA2 samples were excluded from *NSF* analysis because of difficulties in aligning the sequencing reads to this complex region. Lastly, we removed samples whose copy number estimates were too large to be realistic (i.e. copy number > 20 for *KANSL1* and > 50 for *NSF*).

#### Determining haplotypes from 17q21.31 genotypes

We were able to associate the β and α duplications with the inversion status of their haplotype background because the β duplication is always associated with the H1 haplotypes and that the α duplication is always associated with the H2 haplotypes. Therefore, it was possible to integrate the inversion genotypes (H1/H1, H1/H2, H2/H2) and copy numbers (based on read depth in sector calls) to assign complex genotypes representing both inversion and *KANSL1* duplication status as described in^27^ **(Fig. S4)**. We then used R to plot the average copy number across the β and α regions, and assigned genotypes based on their inversion status and copy number. The samples clustered into duplication genotypes with the exception of a handful of samples which did not match their inversion calls. In cases where multiple interpretations of duplication status are possible, the more probable assignment is listed (e.g., H1D/H1D vs. H1/H1DD we chose the former because H1.B3 is very rare). Since there is more diversity in *NSF* copy numbers compared to *KANSL1* especially on the H1.β1 background, we were unable to phase *NSF* duplications and only provide diploid copy number estimates. Complex genotypes for the historical short-read genomes were obtained from Irving-Pease^27^.

#### Sliding window PCAs reveal regions of double recombination

In order to determine recombination or gene conversion events within the inversion locus and extended LD of 100kb, PC1 values were computed in non-overlapping 10kb windows for all individuals using SNPRelate^36^. Tag SNP based inversion genotyping revealed three clusters: H1 homozygous (bottom), H2 homozygous (top), and H1/H2 heterozygous (middle). We computed the mean, median, and multiple quantile ranges of PC1 for individuals of each inversion genotype in every window. To identify noisy or ambiguous positions, we evaluated whether the variance for different genotypes overlapped at each position. Positions with overlapping variance were flagged and excluded from downstream interpretation. The region of interest was further refined to non-duplicated regions except for the unique *KANSL1*. To refine outlier classification, we focused on runs of at least two adjacent outlier windows within the same sample. For these regions, we reassigned the genotype based on directional deviation of PC1: H1H2 genotypes with PC1 below the genotype’s 5th percentile were reassigned to H1H1, while those above the 95th percentile were reassigned to H2H2. Homozygous genotypes were reassigned to H1H2 if their PC1 values deviated substantially in either the positive or negative direction respectively. Further manual curation allowed us to determine single recombinant events and double recombinant events (**Table S4**).

## ACKNOWLEDGEMENTS

We thank R Nielsen, BM Henn, P Moorjani, CT Miller, JM Vazquez and J Chin for helpful discussion. This work was supported by NIH National Institute of General Medicine award R35GM142916 to PHS and NIH National Human Genome Research Institute award R01HG013017 to PHS and a Leakey Foundation award to SS. This manuscript is the result of funding in whole or in part by the National Institutes of Health (NIH). It is subject to the NIH Public Access Policy. Through acceptance of this federal funding, NIH has been given a right to make this manuscript publicly available in PubMed Central upon the Official Date of Publication, as defined by NIH. The content is solely the responsibility of the authors and does not necessarily represent the official views of the National Institutes of Health.

## AUTHOR CONTRIBUTIONS

PHS conceived of the study. SS, RNL, SF, JLR, and PHS performed all analyses and wrote the manuscript. MWM and ANK provided primate cell lines and tissue culture. VB, SGMM, SG, and BH provided data.

## DATA AND CODE AVAILABILITY

All data used in this project are publicly available and described in the ‘Datasets’ section of the Methods and at the following github link https://github.com/sudmantlab/17q21.31_SV_and_evolution which is archived in Zenodo: https://doi.org/10.5281/zenodo.16883463 ^37^. Copy numbers, genotypes, and structural haplotypes can be found in the Supplementary tables. The HPRC data can be obtained at: https://humanpangenome.org/data/. The HGSVC data can be obtained at: https://www.hgsvc.org/resources, The 1KG and HGDP data can be obtained at: https://www.internationalgenome.org/data/. The SGDP data can be obtained at: https://www.simonsfoundation.org/simons-genome-diversity-project/. The ancient data are available on the European Nucleotide Archive under the accession codes PRJEB64656 and PRJEB50857. The unpublished data for bonobos, chimpanzees and gorillas can be obtained at: https://github.com/sudmantlab/17q21.31_SV_and_evolution. The GenomeArk long read haplotype resolved assemblies for primates can be obtained at: https://www.genomeark.org/

## FIGURE TITLES AND LEGENDS

**Supplementary Figure 1 – 24 distinct 17q21.31 structural haplotypes from human long read genome assemblies:** Schematic of 24 unique structural haplotypes found within 210 long-read haplotype-resolved assemblies. Each row represents a unique structural haplotype. Structural haplotypes are arranged using hierarchical clustering based on jaccard distance, with tip sizes scaled by the number of assemblies sharing each haplotype. The dashed red line indicates where clades were collapsed into a single representative structure as shown in **Figure 1B**.

**Supplementary Figure 2 - Comparative gene organization and segmental duplications at the 17q21.31 locus in humans and chimpanzees:** (Top) Gene names and orientation across the human reference (hg38) 17q21.31 inversion locus and extended distal flanking sequence. The black rectangle denotes the inversion breakpoints of the 17q21.31 locus; orange and yellow rectangles denote the segmental duplications; protein coding genes are shown in blue; non-coding transcripts are shown in green; and pseudogenes are shown in purple. (Bottom) Orthologous inversion in chimpanzees shows a larger inversion region (by size) and additional genes in the chimpanzee region that are not shown in the inversion region in humans.

**Supplementary Figure 3 – Inversion genotype assignments mapped atop copy numbers in the β and α regions:** Each point represents an individual with the x-axis denoting the normalized read depth across β and the y-axis denoting the normalized read depth across α. Points are colored according to inversion genotype, assigned using 1271 tag SNPs. Jitter was applied to both the x and y axes to improve visualization of overlapping points.

**Supplementary Figure 4 – Table assigning complex genotype status based on α and β regions:** Cartesian coordinates (x,y) represent complex genotype assignments with numbers indicating the average copy number as discrete values. The value before the comma corresponds to region β, and the value after the comma corresponds to region α. Red cells denote ambiguous cases which can only be distinguished based on 1271 tag SNPs that are required to differentiate between inversion status. Blue cells represent the most conservative assignment which was used in ambiguous cases within the direct or inverted haplotypes.

**Supplementary Figure 5 – Complex genotype at the *KANSL1* region mapped atop copy numbers in the β and α regions:** Each point represents an individual with the x-axis denoting the normalized read depth across β and the y-axis denoting the normalized read depth across α. Points are colored according to complex genotype status, assigned based on the *KANSL1* region using the criteria described in Supplementary Figures 2 and 3. Jitter was applied to both the x and y axes to improve visualization of overlapping points.

**Supplementary Figure 6 – Read depth across the 17q21.31 locus split by CNV assigned by *KANSL1* calls:** Each line represents an individual colored and grouped by complex genotype. The x-axis denotes genome position, and the y-axis shows the average normalized read depth calculated in sliding windows of 1000 bp. Grouping by complex genotype enables visual comparison of CNV patterns across complex genotypes. Horizontal grey lines at the bottom of each plot indicate the positions of the α and β regions, highlighting the differences in read depth between complex genotypes. This visualization complements the discrete copy number genotype assignments shown in Supplementary Figures 3 and 4, illustrating the underlying read depth variation that defines α and β.

## TABLES

**Supplementary Table 1** - Inversion tagging SNPs

**Supplementary Table 2** - Frequencies of H1.β1, H1.β2, H1.β3, H2.α1, and H2.α2 in 1000 Genomes, HGDP, SGDP and CAAPA2 samples

**Supplementary Table 3** - Frequencies of diploid *NSF* copy numbers in 1000 Genomes, HGDP, SGDP and CAAPA2 samples

**Supplementary Table 4** - Recombinant individuals and recombinant breakpoints in 1000 Genomes, HGDP, and SGDP samples

**Supplementary Table 5** - Frequencies of H1.β1, H1.β2, H1.β3, H2.α1, and H2.α2 in Ancient European Genomes

